# Reexamining the evolutionary history of the mammalian medial pterygoid muscle

**DOI:** 10.1101/2025.08.19.671132

**Authors:** Julia A. Schultz, Lucas N. Weaver, Kai R. K. Jäger, David M. Grossnickle

## Abstract

In non-mammalian synapsids, feeding and hearing are closely linked because some jaw bones are involved in both functions. The evolutionary decoupling of these two systems in early mammals likely catalyzed greater specialization of feeding and hearing. Although fossil osteological changes during this process are well documented, the corresponding evolutionary changes to soft tissue anatomy are less certain. The medial pterygoid muscle is a jaw adductor that is central to this evolutionary transition because in many fossil lineages it inserted near or possibly on jaw bones involved in both feeding and hearing. In therians (placentals and marsupials), the medial pterygoid muscle develops medial to Meckel’s cartilage and inserts on the mandibular angular process. Similarly, non-mammalian cynodonts are often reconstructed with a medial pterygoid muscle passing medial to the ossified Meckel’s cartilage, inserting on the dentary ‘angular’ (i.e., pseudangular) process. Thus, the traditional interpretation is that the medial pterygoid remained medial to Meckel’s cartilage through the evolutionary transitions from early cynodonts to therians. Here we highlight issues with that interpretation: the medial pterygoid muscle inserts lateral (not medial) to Meckel’s cartilage in monotremes and, presumably, early mammal groups (e.g., spalacotherioids) that lacked an angular process. This suggests at least two possible explanatory hypotheses: 1) the medial pterygoid muscle is evolutionarily labile, shifting in position relative to Meckel’s cartilage multiple times or 2) the medial pterygoid muscle did not insert on the pseudangular process of non-mammalian cynodonts and instead inserted on the mandibular medial ridge, dorsal to Meckel’s cartilage. We advocate for the latter hypothesis, proposed by Patterson (1956), which suggests that the medial pterygoid did not shift medial to Meckel’s cartilage until the complete separation of the ear and jaw in cladotherians (therians and close relatives), with the shift in position possibly triggering the evolution of the therian angular process as an insertion site. Patterson’s hypothesis is in line with a growing body of evidence that indicate concomitant evolutionary changes of muscles, ears, and jaws at the cladotherian node were important catalysts for the evolution of hearing and feeding specializations in extant mammals.

## INTRODUCTION

Transformation of the jaw bones of non-mammalian synapsids to middle-ear ossicles of mammals is among the best-documented evolutionary transitions in the fossil record, and it likely catalyzed improved feeding and hearing abilities in early Mammalia (Patterson 1956, Rosowski 1992, Luo 2011, Anthwal et al. 2013, Maier and Ruf 2016, Luo et al. 2016, Grossnickle 2017, Bhullar et al. 2019, Luo and Manley 2020, Mao et al. 2020, Mao and Meng 2020, Grossnickle et al. 2022, Schultz 2025). Concomitant with those osteological changes—shrinking, repositioning, and ultimately detaching of bony elements in the jaw into the middle-ear cavity of the skull—were changes in size, position, and direction of contraction of the muscles that facilitated precise tooth-to-tooth occlusion and complex chewing (Fig. 1; e.g., Crompton 1995, Luo and Manley 2020, Grossnickle et al. 2022). The jaw adductor (or elevator) muscles facilitate mammalian chewing, and they include the temporalis, masseter, and pterygoid muscles. Changes to the temporalis and masseter muscles are well traceable through synapsid (mammalian total group) evolution because they are relatively large and their positions are often associated with unambiguous bony features of the synapsid skull, such as the sagittal crest, zygomatic arch, and masseteric fossa (Crompton 1963, DeMar and Barghusen 1972, Bramble 1978, Lautenschlager et al. 2017). The evolutionary history of the medial pterygoid muscle (musculus pterygoideus internus or internal pterygoid), however, is less clear. This includes uncertainty about when in the evolutionary history of synapsids the pterygoideus muscle gave rise to the medial and lateral pterygoid muscles.

**Figure 1.**
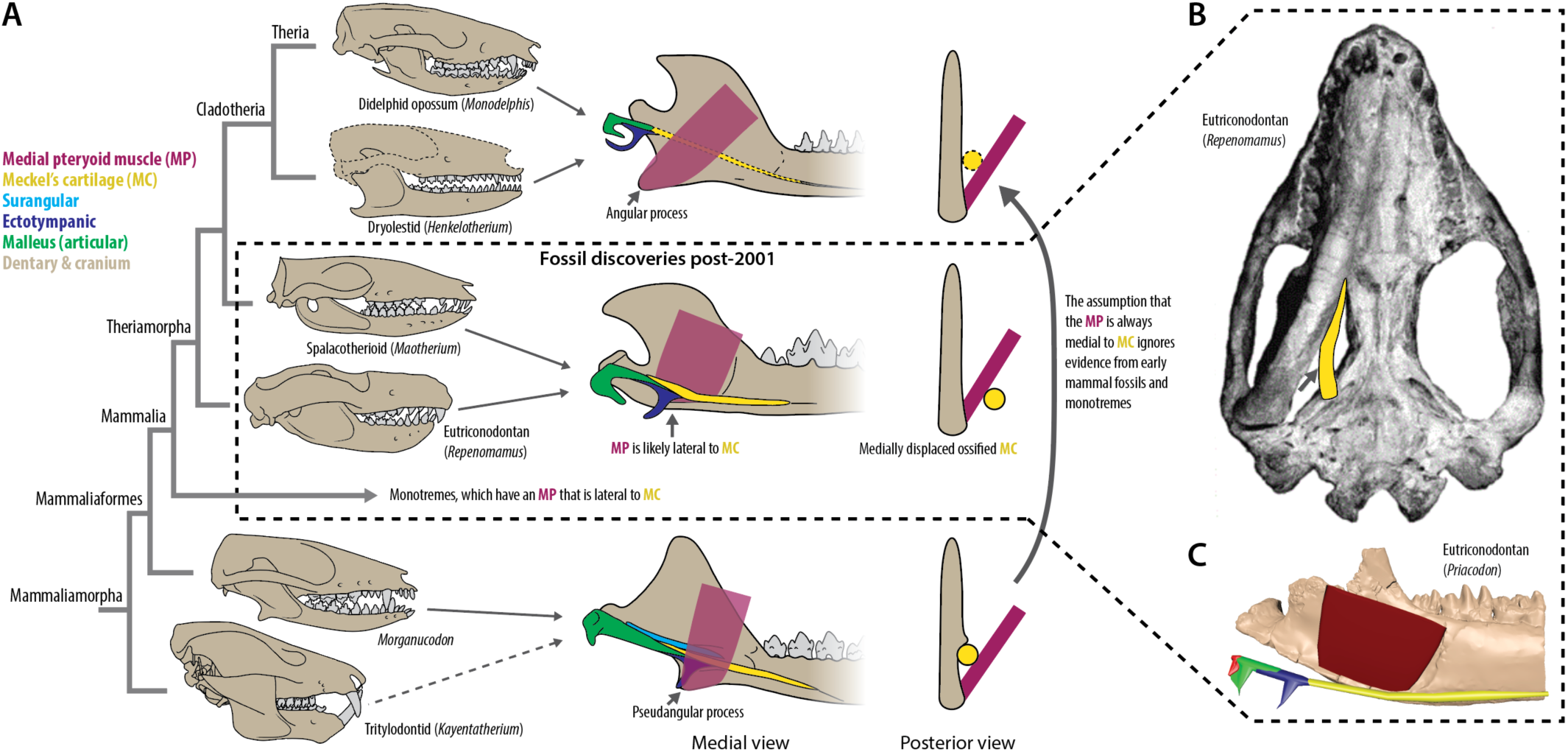
***A***, Summary of the anatomical relationship of the medial pterygoid muscle (MP) and Meckel’s cartilage (MC) during the early evolution of mammals. Medial pterygoid muscle positions are approximate reconstructions. Meckel’s cartilage of cladotherians is illustrated with a dashed line because it either does not ossify or, if ossified, is not medially displaced like those of eutriconodontans and spalacotherioids (e.g., Wang et al. 2022). Early crown mammal such as spalacotherioids and eutriconodontans have medially displaced ossified Meckel’s cartilages (or ‘prearticulars’; Kermack et al. 1973, Wang et al. 2001, Meng et al. 2003, Martin et al. 2015, Mao et al. 2020). Jaw images are of the left hemimandible in medial and posterior view. ***B***, Ventral view of the skull of *Repenomamus* (IVPP V12549), modified from Wang et al. (2003), showing the medial separation of ossified Meckel’s cartilage from the dentary. ***C***, Reconstruction of the ventral portions of the medial and lateral pterygoid muscles in *Priacodon*.

The pterygoideus musculature is linked to the evolution of both the mammalian masticatory and auditory systems because, ancestrally, it likely attached to the postdentary bones of early cynodonts that were involved in both feeding and hearing prior to the separation of the mammalian middle ear (Fig. 1) (e.g., Parrington 1946; Crompton 1963). In extant therians, the medial pterygoid muscle contributes to several types of jaw movements, including yaw (rotation around a dorsoventral axis), roll (rotation around an anteroposterior axis), and lateral translation (Herring and Scapino 1973, Hiiemae and Crompton 1985, Weijs 1980, Weijs 1994, Crompton 1995, Herring 2003, Crompton et al. 2010, Grossnickle 2017, Bhullar et al. 2019, Grossnickle et al. 2022). Prior to the evolution of Cladotheria (therians and their close relatives), however, the pterygoid muscles may have primarily assisted with stabilization of the jaw during mastication rather than facilitating rotational movements (Crompton 1995) because most synapsids likely relied on relatively simple orthal or palinal jaw motions (i.e., with very limited or no yaw, roll, or transverse movements; Grossnickle et al. 2022). In early cladotherians, the medial pterygoid muscle was likely coopted to facilitate more complex anterior and transverse occlusal movements (e.g., Grossnickle et al. 2022 and citations therein), concomitant with the evolution of a posteriorly positioned angular process (Grossnickle 2017) on which the medial surface served as an insertion site for the medial pterygoid muscle.

The medial and lateral pterygoid muscles likely derived from the sauropsid musculus pterygoideus (termed pterygoideus muscle hereafter; Romer and Price 1940, Watson 1953); however, others have proposed that the pterygoideus muscle is homoplastic among tetrapods (Iordansky 2010), and it may have also given rise to additional mammalian muscles beyond the medial and lateral pterygoids, such as the tensor veli palatini and tensor tympani (Barghusen 1986, Crompton et al. 2018). Further uncertainties about the evolutionary history of the pterygoid muscles are related to the phylogenetic appearance of the split of the single pterygoideus muscle into multiple pterygoid muscles, and where those muscles inserted on the jaw. Previous studies alternatively proposed that the medial and lateral pterygoids: (i) arose in early synapsids from a muscle complex (anterior and posterior pterygoideus of Squamata) attaching to the angular bone (Barghusen 1973, King 1981), (ii) evolved in early therapsids from the pterygoideus muscle and attached to the articular/ossified Meckel’s cartilage (e.g., synonymous with the term ‘prearticular’ following Luo and Manley 2020) (Parrington 1955, Crompton 1963, Lautenschlager et al. 2017), or (iii) evolved relatively later in synapsid history (likely in late ‘cynodonts’ but possibly as late as Cladotheria) from the pterygoideus muscle and attached to the dentary (Patterson 1956, Bramble 1978, Barghusen 1986, Crompton 1995). Just like with the insertion sites, there is uncertainty and debate on the cranial origin sites of the pterygoid muscles of synapsids (e.g., Barghusen 1973, Lautenschlager et al. 2017, Crompton 2018). However, compared to jaw elements, the dearth of well-preserved basicranial material in the fossil record makes questions concerning the origin sites even more challenging to examine and unlikely to be resolved at this time.

Most relevant to the evolution of the mammalian chewing and hearing systems is the evolution of pterygoid musculature among cynodonts. There is a general consensus that the pterygoideus musculature shifted from inserting on the postdentary bones to the dentary (see Lautenschlager et al. 2017 for a review) as the postdentary bones reduce in size in the lineage leading from early cynodonts (e.g., *Thrinaxodon*) to mammaliaforms (Crompton 1963A, Allin and Hopson 1992, Sidor 2001). Nonetheless, it is unclear whether that shift occurred early in the evolutionary history of cynodonts (e.g., Crompton 1963B, Kemp 1972), later in the evolutionary history of mammaliamorphans (Bramble 1978, Crompton 1995, Lautenschlager et al. 2017), or among crown mammals (Patterson 1956).

The posteroventral process of the non-therian cynodont dentary—alternatively referred to as the ‘angular process’ or ‘pseudangular process’ by different authors (and hereafter we refer to it as the pseudangular process)—is central to these debates. The pseudangular process first appears among eucynodontian cynodonts and is also present on the dentaries of non-mammalian mammaliaforms. Because the pseudangular process is roughly similar in position and shape to the therian angular process (with the only major difference being the more anterior position of the pseudangular process; Grossnickle 2017), it is often hypothesized that the mammaliaform and therian angular processes are homologous (e.g., Rougier et al. 2015, Abdala and Damiani 2004). Because the medial pterygoid attaches to the medial side of the angular process in extant therians, the insertion site of the medial pterygoid in that evolutionary scenario remained on the ‘angular’ process from early eucynodontians through therians (Adams 1918, Simpson 1928, Crompton 1963, Kemp 1972, Lautenschlager et al. 2017). That the mammal-like division of the pterygoideus muscle into the medial and lateral pterygoid muscles, with the medial pterygoid muscle inserting on the dentary, is generally assumed to have occurred by the origin of Mammaliaformes (e.g., Kermack and Mussett 1981) or potentially earlier among Eucynodontia (Lautenschlager et al. 2017) supports that interpretation. Consequently, the early mammaliaform medial pterygoid muscle, originating on the basicranium, is commonly reconstructed as passing medial to Meckel’s cartilage and the postdentary bones to insert on the ‘angular’ process (e.g., see the *Morganucodon* reconstruction in Figure 1 and Kermack and Mussett 1981). Developmental data from extant therians further support that view—the medial pterygoid muscle also inserts medial to Meckel’s cartilage early in the ontogenetic development of extant therians (Brocklehurst et al. 2015, Sánchez-Villagra and Forasiepi 2017).

Nonetheless, the evolutionary scenario that the medial pterygoid passes medial to Meckel’s from early mammaliaforms to therians is complicated by recent fossil discoveries and phylogenetic analyses demonstrating that several major early mammal clades phylogenetically intermediate to early mammaliaforms and cladotherians (e.g., eutriconodontans, spalacotherioids, multituberculates) lack an angular process (Fig. 1A; Krause et al. 2014, Krause et al. 2020, Rougier et al. 2011, Rougier et al. 2012, Huttenlocker et al. 2017, Luo et al. 2017). These findings suggest that there is an evolutionarily intermediate jaw morphology that lacks the classically invoked insertion location for the medial pterygoid muscle (Fig. 1; Patterson 1956, Patterson and Olson 1961, Grossnickle 2017, Grossnickle 2020, Grossnickle et al. 2022). Thus, we support the interpretation that the pseudangular process is unlikely to be homologous with the therian angular process that evolved at Cladotheria (Patterson 1956, Jenkins et al. 1983, Gow 1986, Grossnickle 2017).

Because some early mammal groups lack an angular process, it is worth reconsidering the evolution of the medial pterygoid and its position relative to Meckel’s cartilage. Although the medial pterygoid muscle is traditionally reconstructed as passing medial to Meckel’s cartilage in non-mammalian synapsids (Fig. 1A), consistent with the early ontogeny of therians (Brocklehurst et al. 2015, Sánchez-Villagra and Forasiepi 2017), Meckel’s cartilage runs very close to the dentary bone in both of those cases (Fig. 1; Fig. 2G-I). Early mammal groups like eutriconodonts and spalacotherioids, however, exhibit a medially inflected Meckel’s cartilage that tethers otherwise-detached middle-ear ossicles to the body of the dentary (Fig. 1B; Wang et al. 2003, Luo et al. 2007, Mao et al. 2020). For the medial pterygoid muscle to insert medial to Meckel’s cartilage in these early mammals, it must have draped around the medially detached middle-ear bones to insert in the deep pterygoid fossa and/or on the ‘pterygoid shelf’ (Sànchez-Villagra and Smith 1997; or ‘pterygoid crest’ of Simpson 1926); such an arrangement is very unlikely because it would have both hindered the vibratory capabilities of the middle-ear complex and left an anomalous wide space empty between the middle-ear bones and the pterygoid fossa. Thus, the medial pterygoid in early mammals was likely positioned lateral to Meckel’s cartilage (Fig. 1B), providing space for the suspended middle-ear elements.

**Figure 2.**
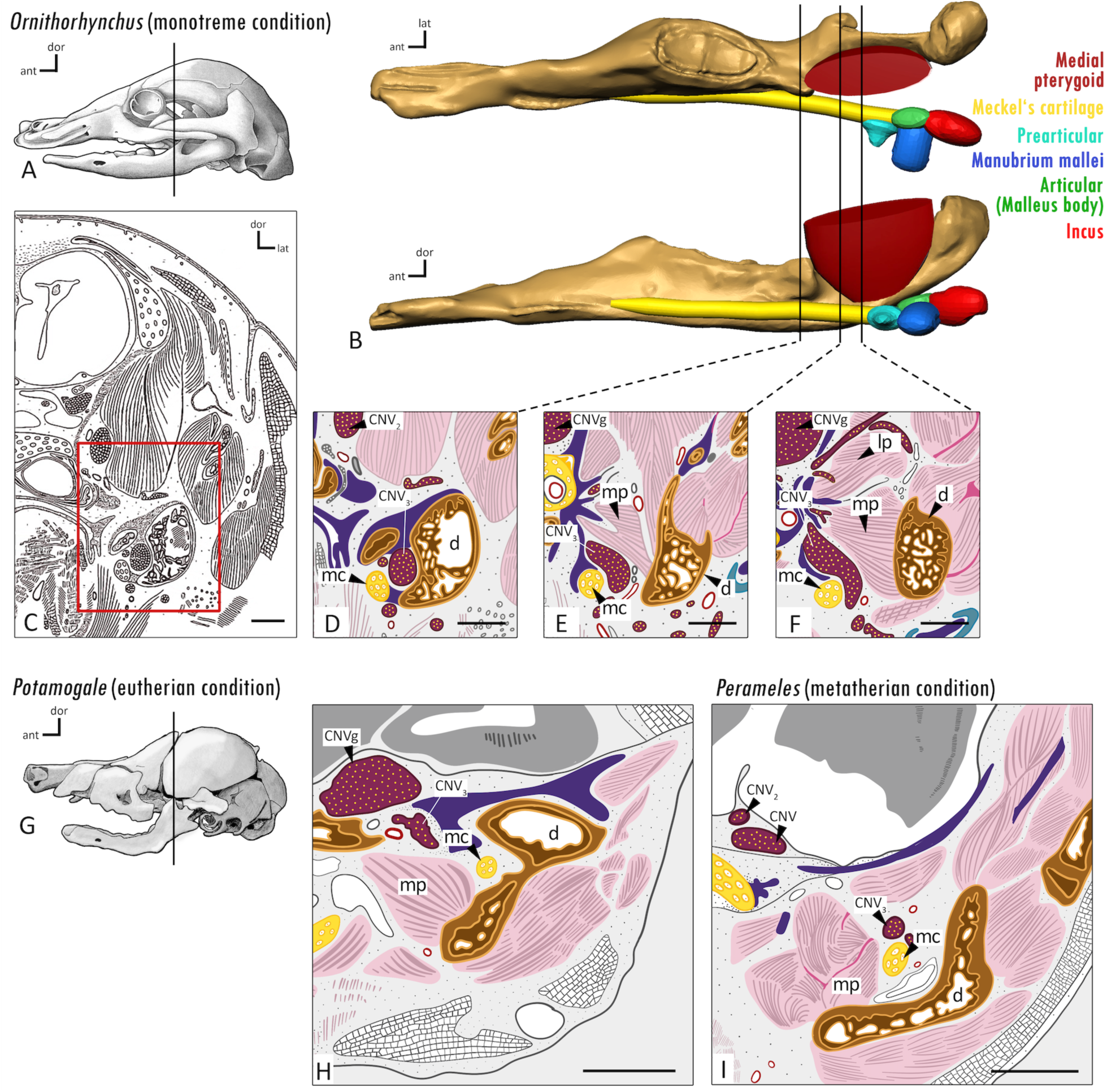
***A*** Chondrocranium reconstruction of a platypus nestling (*Ornithorhynchus anatinus*, 70 mm crown-rump-length) modified from Zeller (1989). The vertical line indicates the orientation of coronal sections in C–F. ***B*** The 3D model of a juvenile platypus lower jaw based on illustrations in Zeller (1989), highlighting the position of the medial pterygoid muscle (pink) between the dentary (brown) and Meckel’s cartilage (yellow). ***C*** Line-drawing of coronal histological section modified from Zeller (1989), with the red box indicating the area shown in D-F. ***D*-*F*** Histological sections modified from Zeller (1989), illustrating the monotreme condition in which the medial pterygoid muscle (mp) passes lateral to Meckel’s cartilage (mc), nested between the Meckel’s cartilage and the dentary bone (d). ***G*** Chondrocranium and dermal bone skeleton of a water shrew fetus (*Potamogale velox*, 105 mm crown-rump-length) modified from Ruf et al. (2020). The vertical line indicates the position of coronal section in H. ***H*** Histological section of a water shrew fetus (*Potamogale velox*, 48 mm crown-rump-length) modified from Brocklehurst et al. (2016), illustrating the eutherian condition in which the medial pterygoid muscle passes medial to Meckel’s cartilage (mc). ***I*** Histological section of a pouch young bandicoot (*Perameles* sp., head length 17.5 mm) modified from Sánchez-Villagra and Forasiepi (2017), illustrating the metatherian condition in which the medial pterygoid muscle passes medial to Meckel’s cartilage. For orientation in *D-F*, *H*, and *I*, the positions of the CNV (fifth cranial nerve or trigeminal), CNV_2_ (maxillary nerve) and CNV_3_ (mandibular nerve), and CNVg (trigeminal ganglion) are indicated. Abbreviations: *ant* anterior *dor* dorsal, *lat* lateral. Scale bars are approximately 1 mm.

In addition to early mammals, monotremes also exhibit a medial pterygoid muscle (or pterygoideus muscle) that passes lateral to Meckel’s cartilage (i.e., the opposite of the therian condition). Although it has been suggested that monotremes lack a ‘true’ pterygoid muscle (Patterson 1956), the presence of the pterygoid (both medial and lateral parts) is reported for *Ornithorhynchus* in multiple studies (e.g., Zeller 1989, Diogo and Powell 2019). Zeller (1989) traced the medial pterygoid in different ontogenetic stages of *Ornithorhynchus* specimens, showing that *Ornithorhynchus* exhibits a medial pterygoid muscle that, early in ontogeny, inserts lateral to Meckel’s cartilage, nestled within the pterygoid fossa with a significant gap between the Meckel’s cartilage and the dentary bone (Fig. 2B, 2D-F); such an arrangement is likely analogous to the muscle and middle-ear arrangement of eutriconodontans and spalacotherioids (Fig. 1). In sum, monotremes and therians appear to have conflicting arrangements of the medial pterygoid muscle relative to the Meckel’s cartilage and middle-ear complex (Fig. 2), mirroring the conflicts raised by the fossil record (Fig. 1).

If the traditional reconstruction for non-mammalian cynodonts is correct—that is, a medial pterygoid muscle inserting on the pseudangular process—then the evolutionary lineage to therians must have included two evolutionary shifts of the medial pterygoid muscle relative to the position of Meckel’s cartilage: one in early mammals, which lack an angular process, and one at Cladotheria when the ‘true’ angular process evolved. These shifts are illustrated in Figure 1A. This raises questions about the accuracy of previous muscle reconstructions and how soft tissues like the medial pterygoid muscle interacted with the detaching middle-ear complex. Answering those questions is crucial for understanding the chewing and hearing mechanisms of early mammals, which lack extant analogs.

Here we review hypotheses about the evolution of the mammalian medial pterygoid muscle in Cynodontia and discuss their implications for homology and function. We propose a novel interpretation of medial pterygoid evolution that builds on a hypothesis proposed by Patterson (1956). Nonetheless, we emphasize that each hypothesis has its merits and limitations, and our aim is to ultimately reignite discussions and provide suggestions for future studies to help resolve this evolutionary conundrum.

## HYPOTHESES ON THE EVOLUTION OF THE MAMMALIAN MEDIAL PTERYGOID MUSCLE

Although there are numerous open questions on the evolution of the medial pterygoid muscle, hereafter we primarily focus on the issue of the medial pterygoid insertion site in the cynodont lineage leading to mammals (Fig. 1). Specifically, we review three alternative hypotheses on the evolution of the mammalian medial pterygoid muscle (Fig. 3) and highlight conflicting lines of evidence derived from what is known from the fossil record and ontogenetic studies of extant mammals, including monotremes. Two of the hypotheses (Simpson Hypothesis and Patterson Hypothesis) were previously proposed but are updated here based on our current understanding of the fossil record, and we introduce a new hypothesis based on phylogenetic parsimony (Fig. 3). We discuss the merits and shortcomings of each hypothesis.

**Figure 3.**
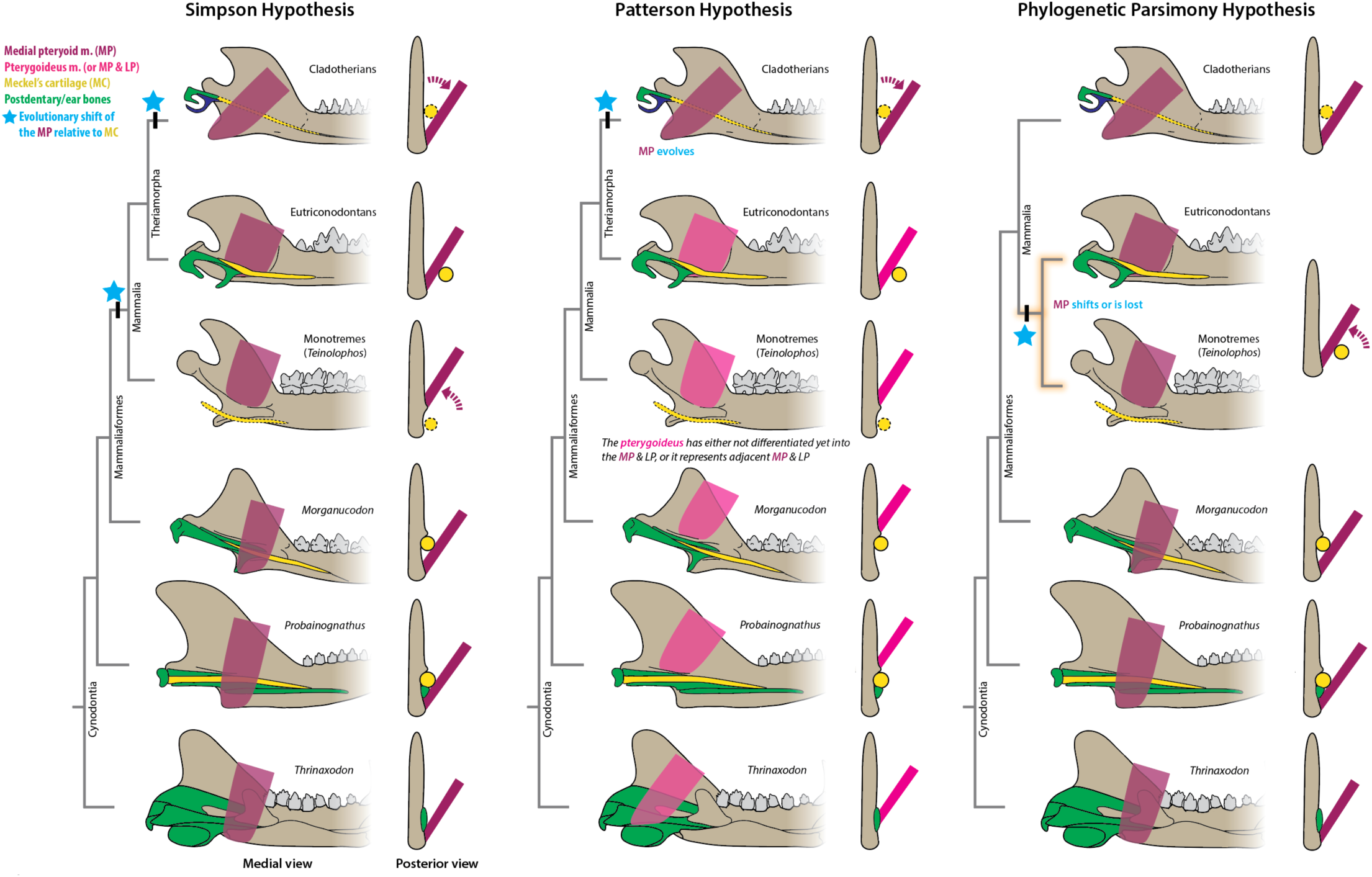
Left hemimandible and medial pterygoid (or pterygoideus) muscle reconstructions in medial and posterior views. The Simpson Hypothesis is the ‘traditional’ reconstruction of the medial pterygoid muscle on the pseudangular process of non-cladotherian cynodonts (Fig. 1; Simpson 1928, Lautenschlager et al. 2017). Following the Patterson Hypothesis, the pterygoideus musculature (pink) may represent an undifferentiated pterygoideus muscle (Patterson 1956), or it may represent separate but adjacent medial pterygoid (MP) and lateral pterygoid (LP) muscles (see the alternative Patterson Hypothesis in the text). The highlighted phylogeny branches in the Phylogenetic Parsimony Hypothesis represent the clade (with spalacotherioids and multituberculates) that would need to be recovered in phylogenetic analyses to support the hypothesis. Blue stars and dashed maroon arrows indicate phylogenetic nodes at which the relative position of the medial pterygoid to Meckel’s cartilage changes. Meckel’s cartilages in early cladotherians (based in part on reconstructions in Allin and Hopson 1992) and early monotremes (represented by *Teinolophos*; Rich et al. 2016) are drawn with dashed lines because those elements are not presented in the fossil record, and in cladotherians it is likely only present early in ontogenetic development. The Meckel’s cartilage position in *Teinolophos* is based on the position of Meckel’s groove (Rich et al. 2016), and additional ear elements are not shown due to uncertainty of their positions.

### 1) Simpson Hypothesis

This hypothesis represents the traditional interpretation of medial pterygoid muscle evolution, which we summarized in the *Introduction*. Its central tenet is that the medial pterygoid muscle inserts on the pseudangular process of the dentary in non-mammalian cynodonts (Figs. 1 and 3; Simpson 1928, Crompton 1963, Lautenschlager 2017). We title this the ‘Simpson Hypothesis’ because, to our knowledge, Simpson (1928) was the first to argue that the mammaliaform pseudangular process is homologous to the therian angular process. In descriptions of the lower jaws of *Morganucodon* (e.g., Mussett and Kermack 1981) and docodontans (e.g., Simpson 1928, Rougier et al. 2015), the pseudangular process is often homologized with the angular process of therian mammals and assumed to be the attachment site of the medial pterygoid muscle in all non-mammalian cynodonts with a developed pseudangular/angular process. In this scenario, the medial pterygoid muscle in non-mammalian cynodonts passes medial to the middle ear bones in the postdentary trough (Fig. 3A). Assuming the pterygoideus muscle (or multiple pterygoid muscles) ancestrally inserted on the angular-articular region of cynodonts (e.g., Lautenschlager et al. 2017), the lateral pterygoid muscle insertion site would have shifted dorsally to the dentary condyle, whereas the medial pterygoid muscle insertion site would have shifted in the opposite direction, ventrally onto the dentary pseudangular process.

#### Conflicting evidence

Some early mammalian groups (i.e., eutriconodontans, spalacotherioids, and multituberculates) lack a pseudangular or angular process and instead exhibit a deep pterygoid fossa on the posteromedial portion of the dentary (Fig. 1). Following the Simpson Hypothesis configuration, the medial pterygoid muscle would therefore have to evolutionarily shift laterally ‘over’ the Meckel’s cartilage and postdentary trough to get from the pseudangular process (medial to the middle ear ossicles and Meckel’s cartilage) to occupy the pterygoid fossa (lateral to the middle ear ossicles and the Meckel’s cartilage), as shown in Figures 1 and 3. From its position lateral to Meckel’s cartilage within the pterygoid fossa, the medial pterygoid muscle would then have to shift again to pass medial to Meckel’s cartilage as is seen among extant therians (and presumably among their close cladotherian relatives; Fig. 3). Thus, the Simpson Hypothesis requires the medial pterygoid muscle to make two evolutionary shifts in position relative to Meckel’s cartilage.

### 2) Patterson Hypothesis

Patterson (1956) posited that the cynodont pterygoideus muscle did not differentiate into the medial and lateral pterygoid muscles until Cladotheria (Fig. 3). He further suggested that the undifferentiated pterygoideus attached to the ‘medial ridge’ (or ‘dentary shelf’; Sidor and Hopson 2017), which is dorsal to Meckel’s cartilage and the postdentary bones of mammaliaforms (Fig. 4), not to the pseudangular process as proposed for the medial pterygoid muscle in the Simpson Hypothesis. In accordance with the Patterson Hypothesis, if a muscle attached to the medial surface of the pseudangular process in non-mammalian mammaliaforms, it was not the medial pterygoid muscle; instead, it may have been jaw-depressor muscles such as the digastric (Bramble 1978) or detrahens mandibulae, the latter of which is unique (as far as we know) to extant monotremes and inserts on the pseudangular process of echidnas (Schulman 1906; referred to as the ‘echidna angle’ by Patterson 1956 and Patterson and Olson 1961) or on the ventral surface of the mandibular body in platypuses (Hopson 1966). As the postdentary bones detached and were medially displaced from the jaw during their transition to mammalian middle-ear ossicles, attachment of the pterygoideus muscles remained along the medial ridge, which is posited to be homologous to the pterygoid shelf or fossa (Fig. 4) in early mammal clades that lack a pseudangular or angular process (i.e., eutricondontans, spalacotherioids, and multituberculates). In this evolutionary scenario, the medial pterygoid muscle shifted from a position dorsal to the postdentary bones to a position lateral to Meckel’s cartilage as the pterygoid shelf shifted ventrally (Figs. 3 and 4), which is similar to the monotreme condition early in ontogeny (Fig. 2B, D-F).

**Figure 4.**
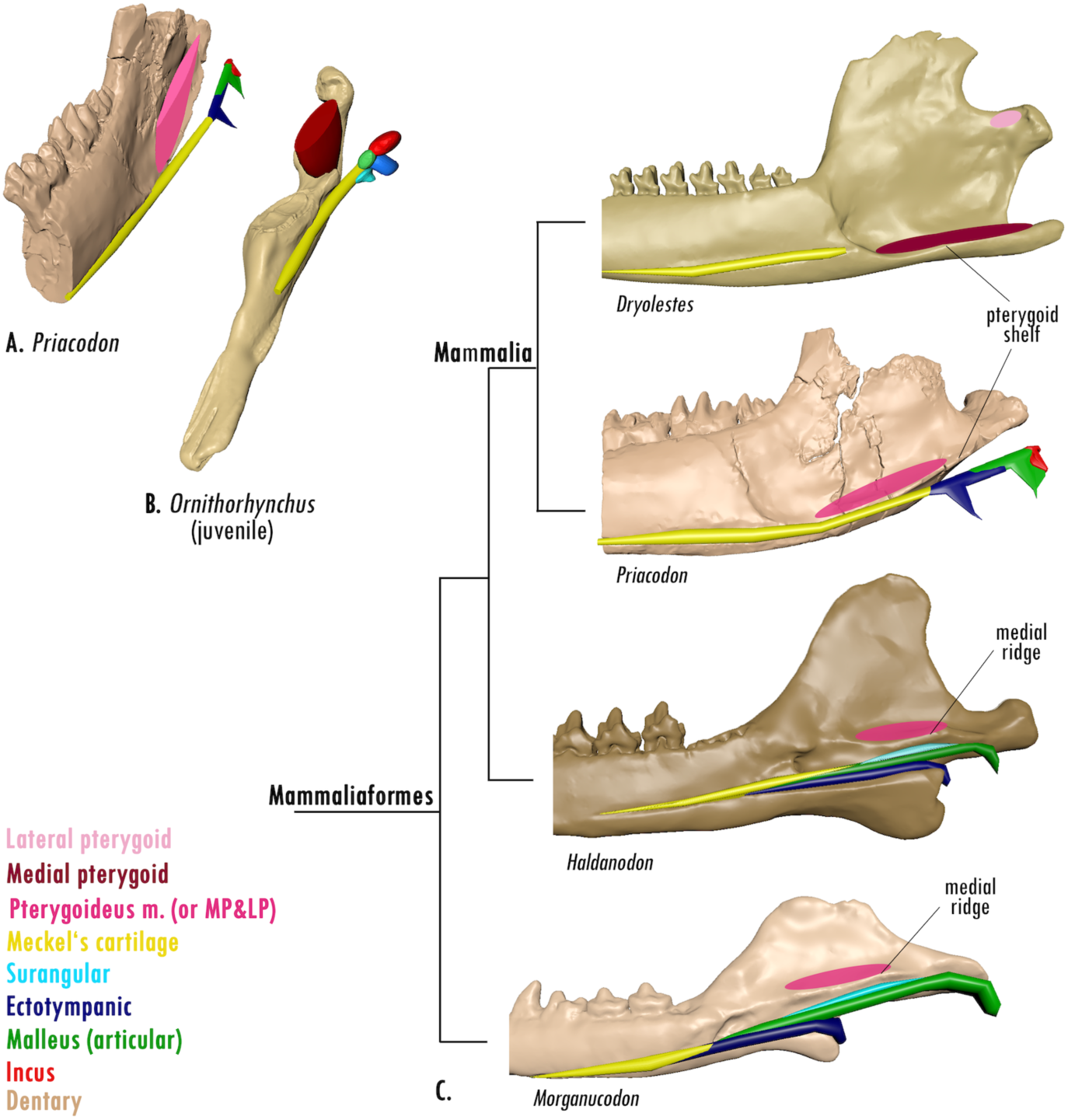
***A***, Posterior part of the lower jaw of the eutriconodontan *Priacodon* with virtual middle ear bones in place to illustrate the lateral position of the pterygoideus musculature (light red) in relation to the middle ear bones. ***B***, Lower jaw of an *Ornithorhynchus* (monotreme) hatchling with the laterally positioned medial pterygoid muscle (dark red) in relation to the middle ear bones. Only the medial pterygoid is reconstructed for *Ornithorhynchus* (see Fig. 2) and not the lateral pterygoid, which is visible in Figure 2F. ***C***, Following the Patterson Hypothesis, we reconstructed the insertion of the pterygoideus musculature (light red), which represents either a single muscle or adjacent medial and lateral pterygoid muscles (because it is unclear if a separation pterygoid muscles existed in those taxa), in lower jaws of the morganucodontan *Morganucodon*, docodontan *Haldanodon*, and eutriconodontan *Priacodon*, and the resulting cladotherian split of both pterygoids (medial [dark red] and lateral [pink]) in *Dryolestes*. Jaws are not to scale.

A central aspect of the Patterson Hypothesis is that the pterygoideus musculature inserted on the medial ridge of non-mammalian synapsids, and not on the pseudangular process (Fig. 4). In this scenario, the medial ridge of non-mammalian synapsids is homologous to the ‘pterygoid shelf’ (medioventral to the pterygoid fossa) of early mammals like eutriconodontans (Fig. 1C). It is worth mentioning that extant mammals often exhibit a ‘endocondylar ridge,’ a structure similar to the medial ridge that likely serves a bite force distributing function (Wilken et al. 2024). However, the endocondylar ridge and medial ridge are likely analogous, not homologous, because many early mammals (e.g., eutriconodontans, multituberculates, spalacotherioids) lack either structure, just like those groups lack an angular process.

An alternative version of the Patterson Hypothesis is that the medial and lateral pterygoid muscles differentiated earlier in synapsid evolution (at least by Mammaliaformes), which is a common view (e.g., Barghusen 1973, Romer and Price 1940, Watson 1953, Crompton 1963, Lautenschlager et al. 2017), and both muscles inserted on the medial ridge (or elsewhere dorsal to Meckel’s cartilage) in non-mammalian mammaliaforms, not on the pseudangular process. In this scenario, the medial pterygoid muscle (rather than the entire pterygoideus muscle like in the original version of the Patterson Hypothesis) evolutionarily drops ventrally between Meckel’s cartilage and the dentary bone in early mammals. This alternative hypothesis is supported by the presence of both a medial pterygoid and lateral pterygoid in monotremes (Fig. 2F; Zeller 1989, Diogo and Powell 2019), which is why we illustrate the medial pterygoid separately from the lateral pterygoid in monotremes (Figs. 2 and 4), except for *Teinolophos* in Figure 3.

#### Conflicting evidence

The original version of Patterson’s hypothesis posits that the pterygoideus does not differentiate into the medial and lateral pterygoid muscles until Cladotheria, which conflicts with interpretations of most other studies on the evolution of mammalian jaw adductor musculature (Simpson 1928, Romer and Price 1940, Watson 1953, Crompton 1963, Barghusen 1973, Kermack and Mussett 1981, Lautenschlager et al. 2017) and the observation of both a medial and lateral pterygoid in monotremes (Fig 2F: Zeller 1989, Diogo and Powell 2019). (Note that the alternative version of the Patterson Hypothesis discussed in the previous paragraph helps to address this conflict.) Further, the Patterson Hypothesis suggests that when the medial pterygoid evolved at Cladotheria it passed medial to Meckel’s cartilage (which is supported by ontogenetic evidence from therians; Fig. 2H, I), whereas the pterygoideus muscle of early mammals passed lateral to Meckel’s cartilage (Fig. 3). In other words, some pterygoideus muscle fibers probably shifted in position relative to Meckel’s cartilage, which would also occur in the scenario presented by the Simpson Hypothesis (Figs. 1 and 3). Nonetheless, adult cladotherians do not maintain the ear-jaw connection via Meckel’s cartilage; the connection is lost early in ontogeny. Thus, there would not have been an ossified Meckel’s cartilage limiting jaw masticatory movements in adults – this may be linked to the evolutionary shift in muscle position (Grossnickle 2017, Grossnickle et al. 2022).

An additional issue with the Patterson Hypothesis is that the superficial masseter muscle is commonly reconstructed to insert on the lateral surface of the pseudangular and angular processes (e.g., Abdala and Damiani 2004, Lautenschlager et al. 2017; but see Patterson and Olson 1961 for an opposing view). If these reconstructions are correct, it supports the argument that pseudangular and angular processes are homologous (e.g., Rougier et al. 2015), conflicting with Patterson’s assertion of non-homology of these structures.

### 3) Phylogenetic Parsimony Hypothesis

This hypothesis, like the Simpson Hypothesis, reconstructs the medial pterygoid muscle as inserting on the pseudangular process of the dentary in non-mammalian cynodonts (Fig. 3; Simpson 1928, Crompton 1963, Lautenschlager et al. 2017). However, it minimizes the number of evolutionary shifts in the position of the medial pterygoid muscle relative to Meckel’s cartilage by collapsing early crown lineages (monotremes, spalacotherioids, eutriconodonts, and multituberculates) into a single clade; thus, it includes only a single shift in medial pterygoid position (or evolutionary loss of the medial pterygoid muscle) at the base of that clade (Fig. 3). If the mammaliaform pseudangular process and cladotherian angular process are homologous (e.g., Rougier et al. 2015), then this hypothesis offers a possible explanation for the lack of an angular process in multiple early mammal lineages; those lineages may form a monophyletic clade and the loss of an angular process (in all groups besides monotremes) may be a derived trait of the clade.

#### Conflicting evidence

The major conflicting line of evidence with the Phylogenetic Parsimony Hypothesis is that it requires a phylogenetic topology that contradicts the phylogenetic hypotheses of multiple independent studies (Meng et al. 2011, Rougier et al. 2012, Krause et al. 2014, Krause et al. 2020, Luo et al. 2015, Luo et al. 2017, Huttenlocker 2018, Hoffmann et al. 2020). To the best of our knowledge, no published phylogenetic analysis has recovered monotremes, spalacotherioids, eutriconodonts, and multituberculates as a clade. Due to the lack of phylogenetic evidence for this hypothesis, we hereafter primarily focus on evaluating the Simpson and Patterson hypotheses. Nevertheless, future studies could test this hypothesis by examining the likelihood of the required tree topology.

## PTERYGOIDEUS MUSCULATURE OF MONOTREMES AND THERIANS

A central observation of this study, which to the best of our knowledge has not been discussed in previous literature, is that therians and monotremes have different positions of the medial pterygoid muscle relative to Meckel’s cartilage during ontogenetic development (Figs. 2 and 3; Zeller 1989, Brocklehurst et al. 2015, Sánchez-Villagra and Forasiepi 2017). Thus, even if fossil evidence is ignored, there is still an issue with the homology of the medial pterygoid muscle among Mammalia. We contend that the Patterson Hypothesis, not the Simpson Hypothesis, aligns more parsimoniously with those embryological studies (Fig. 3). That is, the Patterson Hypothesis does not require the insertion site of the pterygoideus musculature to make an evolutionary shift relative to Meckel’s cartilage in early mammals, whereas the Simpson Hypothesis does (Fig. 3). Following the Patterson Hypothesis, if the pterygoideus musculature inserted on the pterygoid shelf and passed lateral to Meckel’s cartilage in early mammals, then the pterygoideus musculature in monotremes may have simply maintained that ancestral state (Fig. 2; Zeller 1989). (Patterson [1956] argued that monotremes lack a medial pterygoid or pterygoideus muscle, but other authors have shown that the muscle is present in monotremes [Toldt 1905, Zeller 1989, Diogo and Powell 2019].) That the therian medial pterygoid muscle, in contrast, inserts medial to Meckel’s cartilage early in ontogeny may be linked to the evolution of the cladotherian angular process, which might in turn have initiated the reposition of the muscle insertion site. In further support of the Patterson Hypothesis, a human embryo diagnosed with Treacher Collins syndrome (mandibulofacial dysostosis) was observed to have a medial pterygoid that passed *lateral* to Meckel’s cartilage and does not insert on the angular process, instead inserting dorsal to ear elements at or near the mylohyoid line (Herring et al. 1979) – this configuration is consistent with the ancestral mammalian condition following the Patterson Hypothesis.

The jaw morphology of the Early Cretaceous fossil monotreme, *Teinolophos* (Rich et al. 2016), also provides support for the Patterson Hypothesis. In *Teinolophos*, Meckel’s groove is described as ventral to a trough for the mandibular branch of the trigeminal nerve (V_3_) (Rich et al. 2016), suggesting that Meckel’s cartilage passed very close to the ventral margin of the mandibular body (Fig. 3). Thus, there would have been very little space on the medial surface of the pseudangular process for insertion of the medial pterygoid muscle, which would have likely interfered with middle ear elements occupying the same space. More likely is that the pterygoideus musculature inserted dorsal to both Meckel’s cartilage and the trough for V_3_, which is how we reconstruct it in Figure 3. That anatomical configuration for *Teinolophos* (and spalacotherioids and eutriconodonts) is consistent with that of extant monotremes (Fig. 2A–F): the insertion site of the medial pterygoid muscle in *Ornithorhynchus* is primarily dorsal to Meckel’s cartilage, with the muscle fibers passing lateral to Meckel’s cartilage (Figs. 1 and 3).

## THE PSEUDANGULAR PROCESS AND JAW DEPRESSOR MUSCLES

We follow the Patterson Hypothesis (Patterson 1956, Patterson and Olson 1961) in contending that the medial face of the cynodont pseudangular process was not the insertion site for the medial pterygoid, but rather jaw-depressor musculature. (Similarly, Patterson and Olson [1961] argued that no part of the masseter muscle inserted on the pseudangular process.) In sauropsids and early synapsids, the depressor mandibulae was the primary jaw-depressor muscle, and it inserted on the articular, the postdentary bone forming the ancestral jaw joint with the quadrate of the cranium. As the secondary—i.e., dentary-squamosal—jaw joint was established during the evolution of cynodonts, however, the articular became increasingly reduced in size (e.g., Sidor 2003). Hopson (1966) hypothesized that reduction of the articular bone resulted in the ancestral depressor mandibulae losing its function, providing an evolutionary impetus for new jaw-depressor musculature to emerge. He hypothesized that both the therian digastric muscle and the monotreme detrahens mandibulae muscle therefore emerged as novel jaw depressors as a consequence of the establishment of the dentary-squamosal jaw joint in cynodonts.

Indeed, consistent with Hopson’s (1966) hypothesis, the pseudangular process first appears among cynodonts in which the articular-quadrate jaw joint gives way to the dentary-squamosal jaw joint (e.g., *Probelesodon*, *Probainognathus*) (Lautenschlager et al. 2017). Hopson (1966) further hypothesized that reduction of the articular resulted in the loss of function of the sauropsid depressor mandibulae muscle, which attached to the retroarticular process and new jaw-depressor (mouth-opening) muscles like the digastric (of therians) and detrahens mandibulae (of monotremes) could emerge. Following Hopson’s (1966) hypothesis, the sauropsid depressor mandibulae muscles became the “pterygoideus complex” in lineages leading to mammals, with the depressor mandibulae itself likely becoming the tensor tympani muscle of mammals. Thus, reduction of the articular bone may have driven the emergence of both novel mammalian jaw-depressors muscles and a new site for their attachment along the posteroventral and medial portion of the dentary (i.e., the pseudangular process). The view that a muscle other than the medial pterygoid inserted on the pseudangular process in non-mammaliaform cynodonts is shared by Bramble (1978) and seemingly supported by additional studies, such as Crompton and Hylander (1986) and Crompton (1995).

The evolutionary emergence of the pseudangular process also had implications for the superficial masseter, which may have inserted on the pseudangular process’s lateral surface (e.g., Abdala and Damiani 2004, Lautenschlager et al. 2017; but see Patterson and Olson 1961 for an opposing view)—whether changes to the masseter musculature of cynodonts also contributed to the emergence of the pseudangular process, or whether that shift occurred after the structure was in place, remains unclear. Lastly, it is worth noting that some researchers suggest that the pterygoideus also originally attached to the articular in early therapsids (Parrington 1955; Crompton 1963; Lautenschlager et al. 2017). If that was the case, this suggests a rather elegant evolutionary scenario in which the arrangement of the mammalian jaw musculature resulted from a cascade of evolutionary changes brought on by the early establishment of the dentary-squamosal jaw joint in cynodonts.

That the pseudangular process was an attachment site for jaw depressor muscles rather than the medial pterygoid muscle (Patterson 1956, Patterson and Olson 1961) is also supported by comparative anatomy and functional morphology. Relative to the cladotherian angular processes, pseudangular processes of early cynodonts and mammaliaforms are anteriorly positioned relative to the jaw joint, rather than posteriorly positioned (Fig. 5; Patterson 1956, Grossnickle 2017), suggesting that they are not homologous and might be associated with different biomechanical functions. Indeed, the pseudangular process may be functionally analogous to a therian ‘second’ angular process, termed the ‘marginal process’ (Toldt 1905, Patterson 1956) that evolved in some therian lineages, including bat-eared foxes (*Otocyon*), sloth bears (*Melursus*), and *Solenodon* (Fig. 5). In these taxa, the marginal process serves as the insertion site for the jaw-depressing digastric muscle, whereas the medial pterygoid inserts on the ‘true’ angular process (Toldt 1905, Allen 1910, Ewer 1973, Wible 2008). Similarly, the pseudangular process of monotremes, including that of the echnida and *Teinolophos*, is the insertion site of the detrahens mandibulae muscle, a jaw depressor analogous to the therian digastric muscle (Fig. 4; Schulman 1906, Hopson 1966). The relatively anterior position of the pseudangular process compared to the angular process may optimize leverage for jaw depression by the posteriorly oriented jaw depressor muscles. Conversely, the more posteriorly positioned cladotherian angular process is posited to be associated with generating rotational jaw movements such as yaw (Grossnickle 2017, Grossnickle 2020) or roll (Bhullar et al. 2019), which both result in transverse movements of molars during occlusion.

**Figure 5.**
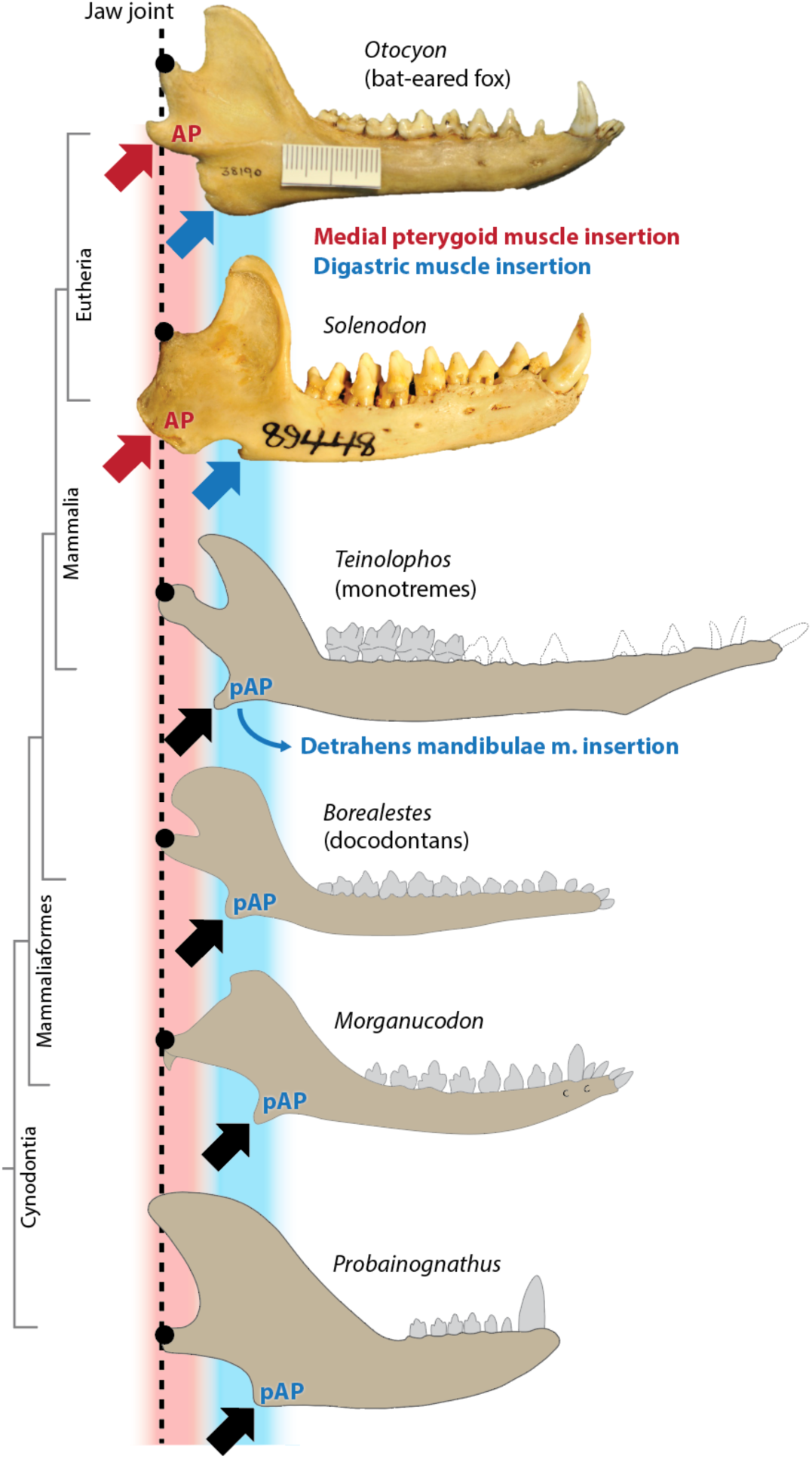
Comparative jaw anatomy of example eutherians that exhibit a ‘marginal process’ (red arrows; Patterson 1956), which is the insertion site for the digastric muscle, and fossil cynodonts. (Note that the eutherian jaws are in lateral view, but the medial pterygoid muscle inserts primarily on the medial surface.) In terms of anatomical position along the length of the jaw, the pseudangular process (pAP) of non-cladotherian cynodonts is more in-line with the therian ‘marginal process’ than the therian angular process (AP; blue arrows). Jaw sizes are scaled roughly based on the jaw-joint-to-molars distance.

Further, the jaw pseudangular process is often described as bent laterally (‘effected angle’; see Schultz et al. 2017 for a summary), which may be due to the pull of the laterally attached superficial masseter lacking a corresponding pull by a medially attached muscle such as the medial pterygoid. In contrast, the angular process of therians, which have a definitive medial pterygoid muscle, is inflected medially in many taxa, such as in most marsupials (e.g., Maier 1987, Sánchez-Villagra and Smith 1997), likely enhancing roll rotation (Grossnickle 2020).

In sum, these observations on functional jaw anatomies support the argument by Patterson (1956) and Patterson and Olson (1961) that (1) the pseudangular and angular processes are not homologous and (2) that the pseudangular process served as an insertion site for a jaw depressor muscle, not the medial pterygoid (Fig. 5), and thus serves a different functional purpose than the angular process.

## EXPANDING ON THE PATTERSON HYPOTHESIS

The Patterson Hypothesis makes two arguments that contradict traditional muscle reconstructions (i.e., the Simpson Hypothesis): (1) the pterygoideus muscle does not give rise to medial and lateral pterygoid muscles until Cladotheria and (2) the pterygoideus muscle inserts on the dentary medial ridge (not the pseudangular process as commonly reconstructed) in non-mammalian mammaliaforms (and possibly earlier taxa), suggesting homology of the medial ridge and medial surface of the therian angular process. In this section, we examine these arguments in greater detail.

The first argument of the Patterson Hypothesis is challenging to assess due to the small size of the lateral pterygoid muscle (making it unlikely for researchers to observe bone scarring or osteological correlates) and the considerable uncertainty on the insertion sites of jaw musculature in non-mammalian synapsids (see the *Introduction*). The therian lateral pterygoid muscle is generally oriented to generate medially or anteriorly directed jaw movements and is often associated with fine-scale jaw movements, such as slight protrusion and/or lateral movement of the jaw in primates (Hylander 2006). But most non-cladotherian synapsid lineages lacked complex chewing cycles that involved transverse movements, instead relying primarily on orthally or posteriorly directed jaw movements (Grossnickle et al. 2022 and sources within), with taxa such as docodontans (with pseudo-tribosphenic molars analogous to therian molars) and the dicynodont *Sangusaurus* (Angielcyzk et al. 2017) being possible exceptions.

We also reiterate that an alternative version of the Patterson Hypothesis is that the medial and lateral pterygoid muscles evolved prior to Cladotheria but that the medial pterygoid never inserted on the pseudangular process. If the ancestral cynodont condition is that both lateral and medial pterygoid muscles are present and lie next to each other in close proximity inserting on the angular-articular region (like has been reconstructed in *Thrinaxodon* and *Probainognathus*; see Lautenschlager et al. 2017 and sources within, Parrington 1955, Barghusen 1968, 1972, Crompton 1963), then the muscles may have remained in close association and both shifted dorsally to the medial ridge. In other words, rather than a single pterygoideus muscle migrating dorsally to the dentary as in the Patterson Hypothesis, it is possible that both the medial and lateral pterygoid muscles shifted dorsally. This contradicts the Simpson Hypothesis in which there is a divided state of the lateral and medial pterygoid muscles in early mammaliaforms, with the lateral pterygoid muscle attaching close to the articular condyle (dorsal of postdentary bones) and the medial pterygoid muscle inserting ventral to the postdentary bones on the pseudangular process.

The second argument of the Patterson Hypothesis—that the pterygoideus muscle inserts on the dentary medial ridge in non-mammalian mammaliaforms—can be assessed by examining osteological evidence and comparing this scenario to that of the Simpson Hypothesis (e.g., Crompton 1963, Lautenschlager 2017). For instance, in favor of the pterygoideus muscle inserting on the dentary medial ridge is that non-mammalian mammaliaforms exhibit a well-developed medial ridge, a bony crest above the postdentary trough (Fig. 4). The medial ridge includes significant surface area (likely greater than the surface area of the medial side of the pseudangular process in most taxa) for muscle attachment. However, it is also worth noting that beyond serving as a muscle insertion site, the medial ridge might serve as the primary load path for distributing bite forces like has been suggested for the endocondylar ridge of therian mammals (Panagiotopoulou et al. 2020; Wilken et al. 2024). Non-mammalian mammaliaforms have complex middle ear elements inside the grooves of the postdentary trough for sound transmission; a pterygoideus that inserts dorsal to the middle ear elements is unlikely to have interfered with sound transmission. In contrast, a medial pterygoid muscle inserting on the pseudangular process (i.e., the Simpson Hypothesis condition; Fig. 3) would pass adjacent to the ear elements, possibly limiting the vibration and thus function of the ear elements.

The Patterson Hypothesis is further supported by our current knowledge of the evolutionary detachment of the middle ear from the dentary. The middle ear ossicles detach posteriorly first while they keep a connection to the lower jaw anterior to the middle ear ossicles via the Meckel’s cartilage (Luo et al. 2007, Mao et al. 2020, Mao and Meng 2020). Following the Patterson Hypothesis, when ear elements are still anteriorly connected to the lower jaw via the Meckel’s cartilage the pterygoideus remains lateral to Meckel’s cartilage (Figs. 1 and 3), even as the pterygoid fossa expands ventrally. For example, in eutriconodonts such as *Yanoconodon*, the pterygoid fossa and the jaw joint are confluent (Luo et al. 2007, Luo 2007) and the pterygoideus may have spread along the entire region, keeping posterior fibers close to the jaw joint, which is where the lateral pterygoid muscle is positioned in extant mammals. In contrast to the Simpson Hypothesis, this evolutionary scenario is possible without a ‘jumping’ of the medial pterygoid muscle over middle ear elements at/near Mammalia (Fig. 3). Luo and Manley (2020) suggest the pterygoid fossa of mammals has ‘replaced’ the non-mammalian mammaliaform postdentary trough, but our interpretation of evidence from monotreme ontogeny and the preserved ossified Meckel’s cartilages of eutriconodontans and spalacotherioids goes further, implying a homology of the non-mammalian mammaliaform attachment area dorsal to the medial ridge and the mammalian pterygoid fossa.

## THE ALLOTHERIAN PERSPECTIVE

One major early mammalian group—Allotheria—has not yet been addressed, due in part to uncertainties surrounding their phylogenetic relationships. Allotheria, if a valid monophyletic group, include haramiyidans (including the Late Triassic-Early Jurassic *Haramiyavia* and *Thomasia* and the Middle Jurassic Euharamiyida), gondwanatherians, and multituberculates (e.g., Butler 2000; Zheng et al. 2013; Mao et al., 2023). However, many authors view ‘Allotheria’ as polyphyletic, with many specific variations in terms of which groups are included/excluded in Allotheria (Butler 2000, Luo et al. 2002, Kielan-Jaworowska et al. 2004, Butler and Hooker 2005, Zhou et al. 2013, Luo et al. 2015b, Krause et al. 2020a, Hoffmann et al. 2020, King and Beck 2020). Disputes surrounding the putative monophyly of Allotheria are centered on interpretations of the middle-ear configuration in haramiyidans. Multituberculata have a fully detached mammalian middle ear (Lillegraven and Hahn 1993, Szalay et al. 1993, Hurum et al. 1996, Rougier et al. 1996). In comparison, some authors contend that all haramiyidans had middle-ear bones still closely attached to the jaw (e.g., Zhou et al. 2013; Luo et al. 2015b), but some euharamiyidans may have exhibited partly attached middle ears still tethered to the jaw by Meckel’s cartilage (i.e., analogous to eutriconodontans and spalacotherioids) and some may have had fully detached definitive mammalian middle ear (e.g., Zheng et al. 2013; Mao et al. 2023). Until the phylogenetic relationships are resolved, interpreting allotherian medial pterygoid muscle evolution remains challenging. Here, we briefly summarize the implications (following the Patterson Hypothesis) under three competing evolutionary scenarios: (1) Allotheria are monophyletic and outside of crown Mammalia, (2) Allotheria are monophyletic and within crown Mammalia, and (3) Allotheria are polyphyletic.

If Allotheria are monophyletic and outside Crown Mammalia, the evolution of the middle ear and medial pterygoid muscle would largely mirror what we have proposed above for non-allotherian mammaliaforms (Fig. 6). Like *Morganucodon*, the coeval *Haramiyavia* had middle-ear bones closely affixed to the jaw in a post-dentary trough (Luo et al. 2015b), whereas, many euharamiyidans had a detached middle ear (e.g., Meng 2014; Mao et al., 2023) or a middle-ear that was tethered to the dentary by Meckel’s cartilage like eutriconodontans and spalacotherioids. Like cladotherians, multituberculates exhibit a fully detached middle ear (e.g., Hurum et al. 1996; Rougier et al. 1996). Following the Patterson Hypothesis in this scenario, the pterygoideus muscle would have attached to the ‘medial ridge’ or ‘pterygoid shelf’ overhanging the postdentary trough in early allotherians like *Haramiyavia* (Luo et al. 2015b; Fig. 6), and it would have then shifted ventrally (and lateral to Meckel’s cartilage) to continue attaching along the prominent pterygoid shelf in euharamiyidans, gondwanatherians, and multituberculates (e.g., Gambaryan and Kielan-Jaworowska 1995, Meng et al. 2014, Krause et al. 2020b). Allotherians did not develop an angular process like that seen in Cladotheria and the medial pterygoid (or pterygoideus) muscle presumably remained lateral to Meckel’s cartilage throughout allotherian evolution. If Allotheria are monophyletic but within Crown Mammalia, the story largely remains the same, with the primary difference that the full sequence of middle ear detachment (from fully attached, to partly detached, to completely detached) occurred independently in allotherians within Crown Mammalia.

**Figure 6.**
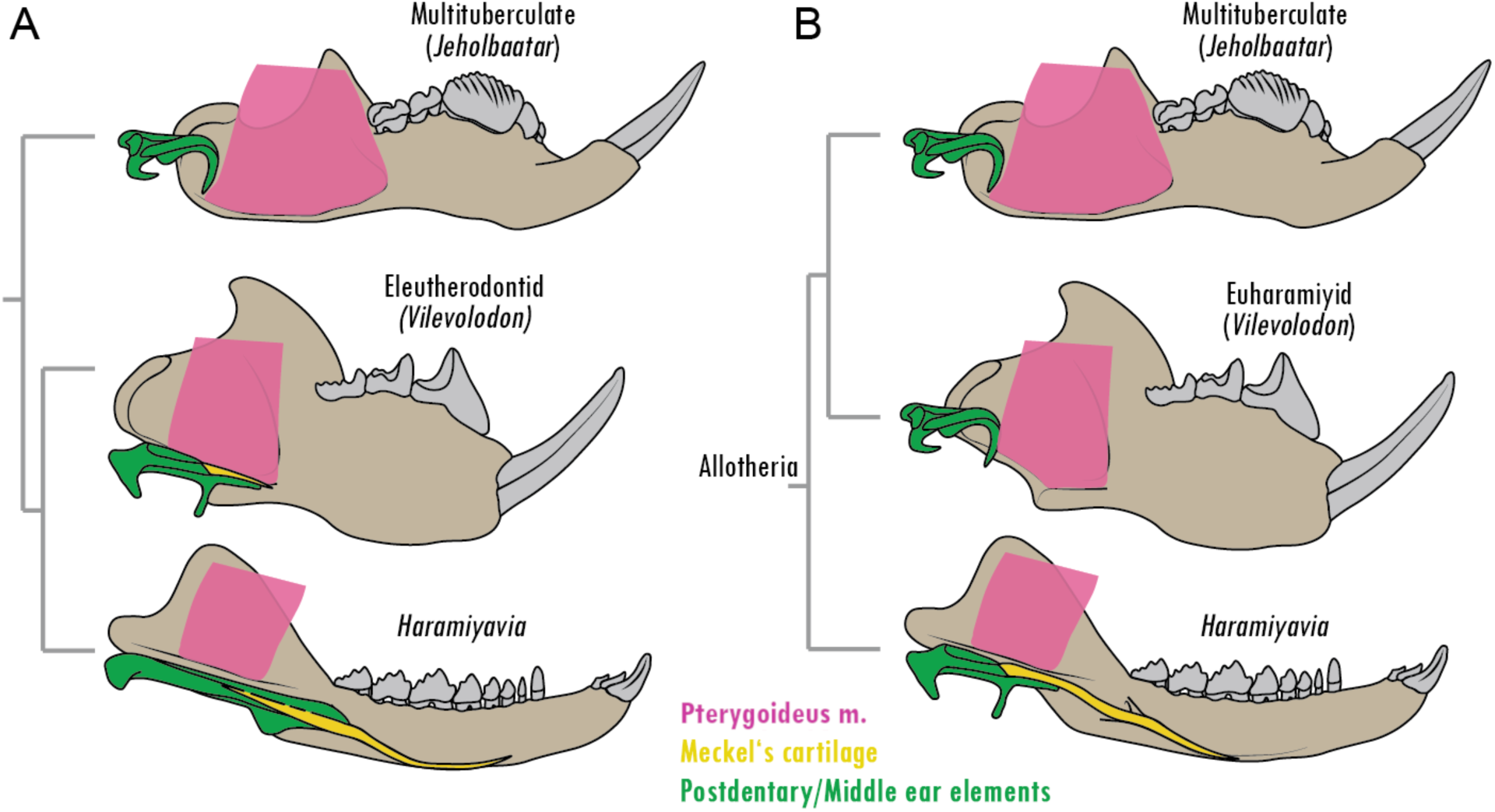
Evolution of the allotherian pterygoid musculature following the Patterson Hypothesis. Whether haramiyidans are non-mammalian mammaliaforms (A; e.g., Luo et al. 2015b) or crown mammals closely related to multituberculates in a monophyletic Allotheria (B; e.g., Zheng et al. 2013), the undifferentiated pterygoid muscle likely attached to the medial ridge or pterygoid shelf above the postdentary elements, either dorsal (as in *Haramiyavia* and, in scenario A, *Vilevolodon*) or lateral (as in *Jeholbaatar* and, in scenario B, *Vilevolodon*) to Meckel’s cartilage.

If Allotheria are polyphyletic and multituberculates are the only crown mammalian group, multituberculates evolved a detached middle ear independently with no obvious evolutionary precursor known from the fossil record. The earliest multituberculate specimens for which the morphology of the posteromedial dentaries can be observed (e.g., Paulchoffatiidae like *Pseudobolodon* Hahn 1969 or *Rugosodon* Yuan et al. 2013), that show no evidence of Meckel’s cartilage (or other post-dentary bones) attaching to the jaw. The medial pterygoid (or pterygoideus) muscle is reconstructed to attach to the pterygoid shelf along the ventral margin of the posteromedial dentary (e.g., Simpson, 1926; Wall and Krause, 1992; Gambaryan and Kielan-Jaworowska, 1995; Fig. 6), but its origin (i.e., medial ridge or pseudangular process) is not clear due to missing evidence in the fossil record leading to Multituberculata.

Relevant to the Patterson Hypothesis, the medial lower jaws of multituberculates do not exhibit obviously different putative insertion locations for a split medial and lateral pterygoid. Instead, the ventromedial portion of the jaw posterior to the molars is simply characterized by an expansive fossa (e.g., Clemens and Kielan-Jaworowska, 1979; Kielan-Jaworowska et al., 2004). Indeed, in previous muscle reconstructions of the multituberculate masticatory apparatus, the medial and lateral pterygoids are either reconstructed as being right next to each other in the pterygoid fossa (with the lateral pterygoid slight posterodorsal to the medial pterygoid; Gambaryan and Kielan-Jaworowska, 1995) or undifferentiated as a single ‘pterygoid’ (Simpson, 1926; Wall and Krause, 1992). Thus, the anatomy of multituberculate jaws in consistent with the Patterson Hypothesis, which assumes the pterygoideus remained a single muscle until the evolution of Cladotheria (Fig. 6).

Unlike nearly all other non-mammalian mammaliaforms, allotherians are characterized by bilateral mastication and exhibit mostly orthal and palinal (posteriorly oriented) chewing strokes (Krause 1982, Jenkins et al. 1997, Butler 2000, Butler and Hooker 2005, Lazzari et al. 2010, Weaver and Wilson 2021, Grossnickle et al. 2022). Thus, whereas the medial pterygoid muscle is essential for facilitating yaw, roll, and translation during the unilateral occlusion of cladotherians, it was likely primarily used for stabilizing the jaws during bilateral occlusion in allotherians (e.g., Butler 2000; Butler and Hooker 2005; Gambaryan and Kielan-Jaworowska 1995). That the pterygoid shelves and fossae of allotherians are very well-developed points to the importance of the pterygoid musculature in general; yet, instead of enhancing transverse movements like cladotherians (e.g., Grossnickle 2017), the medial pterygoid (or pterygoideus) muscle probably helped prevent deviations from the horizontal plane, maintaining the precise occlusion of interlocking longitudinal cusp rows during the palinally directed power stroke that characterizes allotherian mastication.

## IMPLICATIONS FOR HEARING AND FEEDING

At the time of Patterson’s (1956) publication, well-preserved fossils of eutriconodontans and spalacotherioids were unknown, and our understanding of early mammalian phylogenetic relationships were more poorly resolved, so it is unclear how he would have interpreted these new revelations. Patterson (1956) did posit, however, that the medial pterygoid muscle evolved concurrently with the evolutionary disconnection of the middle ear bones of the jaw, talonid shelf of molars, and cladotherian angular process. Following the Patterson Hypothesis, this suggests that specializations of the masticatory and hearing apparatuses evolved in concert with evolution of the pterygoideus musculature. See Grossnickle et al. (2022) for an expanded discussion on how these evolutionary changes may be linked to functional changes of the masticatory apparatus.

The postdentary trough of early mammaliaforms, containing bony and cartilaginous elements and responsible for sound transmission to the inner ear, naturally requires freedom of movement for transmitting vibrations from one element to the other. Any jaw movement (not necessarily chewing) probably interfered with such a fine-tuned system. In particular, soft tissue structures, such as muscles and tendons, as reconstructed with the medial pterygoid muscle attached to the pseudangular process (e.g., see the Simpson Hypothesis in Figure 3), may have hindered transmission process by wrapping closely around the elements preventing mobility. The alternative placement advocated here (Patterson Hypothesis; Fig. 3) of muscle and tendon above the postdentary trough allows for more freedom of movement.

Following the Patterson Hypothesis, once in the pterygoid fossa, the pterygoideus muscle passes dorsal to (in early mammaliaforms) or lateral to (in early mammals) the middle ear elements (Fig. 4). A pterygoideus muscle dorsal or lateral to the middle ear elements prevents possible close contact between the contracting muscle and ear elements. In contrast, following the Simpson Hypothesis scenario in non-mammalian synapsids (Fig. 3), the medial pterygoid muscle would drape medio-ventrally around Meckel’s cartilage, and the contracting muscle would likely interfere with the mobility of the ossicles and limit the transmission of vibrations. Later, in early cladotherians, the ear completely separates from the jaw, which may have allowed the jaw joint to shift dorsally (e.g., Grossnickle 2017) and provide space for the medial pterygoid insertion site to shift to a more posteroventral position on the jaw, likely triggering the evolution of the angular process. As a result, cladotherians reflect the modern mammalian configuration of the medial and lateral pterygoid muscles (Fig. 4). The concurrent evolutionary change to cladotherian molars, namely the origin of a talonid shelf that suggests the evolution of novel medially directed occlusal movements (Schultz and Martin 2014, Grossnickle 2017, Grossnickle et al. 2022), further suggests that the new jaw and muscle morphologies were associated with a fundamental shift in chewing mechanics in cladotherians.

## CONCLUSIONS

There is considerable uncertainty about the position of the medial pterygoid muscle insertion site throughout the evolution of synapsids. For instance, there is a lack of a consensus on the cynodont pterygoideus configuration (Crompton 1993, Bramble 1978, Barghusen 1986, Allin and Hopson 1992, Crompton 1995, Lautenschlager et al. 2017), which is necessary to understand the evolutionary steps that lead to the mammalian condition of the pterygoid muscles. Newly interpreted evidence from fossils and developmental investigations, in addition to the vetting of three competing hypotheses (Fig. 3), highlight the difficulties of understanding the muscular construction of jaw adductors in the history of mammals, but also allow for new perspectives in regard of the evolution of the mammalian hearing and feeding system.

We favor Patterson’s (1956) hypothesis that medial pterygoid muscle did not insert on the pseudangular process in mammaliaforms, based on information derived from recent fossil findings (Figs. 1, 3–6), monotreme ontogeny (Fig. 2), and comparative anatomy of mammalian jaws (Fig. 5). Mammalian evolution may have involved migration of the pterygoideus muscle insertion site from the articular and/or ossified Meckel’s cartilage (i.e., prearticular) in early cynodonts to the dentary’s medial ridge above the postdentary trough in early mammaliaforms (Fig. 4), potentially in response to articular reduction and the establishment of the dentary-squamosal jaw joint (Hopson, 1966). Patterson (1956) argued that the pterygoideus does not give rise to the medial and lateral pterygoid muscles until Cladotheria, but it is also possible that the muscles were present in earlier lineages but both maintained insertion sites dorsal and/or lateral to Meckel’s cartilage. The Patterson Hypothesis indicates homology of the non-mammalian mammaliaform medial ridge and mammalian pterygoid fossa due to the continued insertion of the pterygoideus on this structure during early mammalian evolution. During the evolutionary separation of the middle ear elements from the jaw, the medial pterygoid muscle likely ‘dropped’ ventrally between the middle ear and dentary bone, occupying space lateral to the ossified Meckel’s cartilage. The monotreme condition is a pterygoid muscle that passes lateral to Meckel’s cartilage (Fig. 2), which is the same configuration reconstructed in early crown mammals (Fig. 1), suggesting that monotremes have maintained the ancestral mammalian state. In further support of the Patterson Hypothesis, we present evidence that the medial or posterior surface of the pseudangular process of non-mammalian mammaliaforms was the insertion site of a jaw depressor muscle (possibly the monotreme detrahens mandibulae or the therian digastric; Fig. 5) rather than the medial pterygoid muscle (as posited by the Simpson Hypothesis and Phylogenetic Parsimony Hypothesis). We believe that the evolutionary scenario of the Patterson Hypothesis is more parsimonious than that suggested by the Simpson Hypothesis, which would require multiple changes in the position of the medial pterygoid muscle relative to Meckel’s cartilage (Figs. 1 and 3).

Nonetheless, we emphasize that none of the hypotheses presented in this study (Fig. 3), including the Patterson Hypothesis, are fully satisfactory because they each require seemingly unlikely evolutionary scenarios. Each hypothesis requires either evolutionary shifts in the position of medial pterygoid muscle relative to Meckel’s cartilage or suggest timings of the origin or loss of the medial pterygoid muscle that are inconsistent with various lines of evidence (e.g., fossil record, ontogeny, phylogenetic parsimony). Thus, we recommend future studies further test competing hypotheses on the evolution of medial pterygoid muscle, making use of various lines of evidence from paleontology, developmental biology, phylogenetics, and biomechanics. In particular, the strengths of competing hypotheses are sensitive to the ancestral position of the medial pterygoid muscle on the postdentary elements in non-mammaliaform cynodonts—a ventral position on the pseudangular process would support the Simpson Hypothesis and a dorsal position on postdentary bones or the medial ridge would support the Patterson Hypothesis. This topic could be investigated in mammaliamorphan cynodonts such as brasilodontids (Rawson et al. 2024), which possess a mix of mammalian and non-mammalian traits. In addition, we emphasize that future studies should not only utilize the ever-growing fossil record but should also examine the ontogeny of soft tissue parts surrounding the developing middle ear in extant mammals, especially monotremes. Finally, we see potential for utilizing biomechanical studies to test evolutionary hypotheses on the medial pterygoid (and additional muscles like the superficial masseter), such as investigating load-bearing patterns on non-mammalian mammaliaform lower jaws by comparing the two different attachment sites (pseudangular process of the Simpson Hypothesis versus medial ridge of the Patterson Hypothesis). Our goal here is to reinvigorate interest and debate on the topic of soft tissue evolution in early mammals, and we hope to spur future research on the topic.

## MATERIALS & METHODS

We produced 3D digital models of fossil jaws of the mammaliaforms *Morganucodon*, *Haldanodon*, the eutriconodont *Priacodon*, and the cladotherian *Dryolestes*, which were then used to virtually reconstruct origin and insertion areas of the pterygoides (or medial pterygoid and lateral pterygoid) muscles. The 3D models were produced from micro-computed tomography (micro-CT) scans. The lower jaws of *Morganucodon* (NHM M84028, Natural History Museum London) and *Priacodon* (LACM 120451, Natural History Museum of Los Angeles) were scanned at the University of Texas High-Resolution X-ray CT facility using the NSI scanner. Scan parameters for *Morganucodon*: Xradia, 60 kV, 10 W, 1 s acquisition time, 1 mm SiO2 filter, voxel size 19.64 µm. Scan parameters for *Priacodon*: Fein focus high power source, 110 kV, 0.15 mA, no filter, voxel size 10.4 μm. For further scan information, see Jäger et al. (2020). The lower jaws of *Haldanodon* (GuiMam 3-80) and *Dryolestes* (GuiMam 3-78) were scanned at the Bonner Insitute of Organismic Biology (BIOB, formerly Institute of Geosciences, Universität Bonn; section Palaeontology) using the v|tome|x s µCT device (GE Sensing & Inspection Technologies GmbH phoenix|x-ray). Scan parameters for *Haldanodon*: 240 tube, 90 kV, 180 µA, 333 ms exposure time, no filter, voxel size 16.49 µm. Scan parameters for *Dryolestes*: 240 tube, 90 kV, 180 µA, 333ms exposure time, no filter, voxel size 24.66 µm.

The 3D model of the *Ornithorhynchus* hatchling lower jaw is a virtually edited polygonal surface model based on the adult surface model of *Ornithorhynchus anatinus* downloaded from MorphoSource (imnh:r:2125; OBJ mesh file, Idaho Museum of Natural History specimen S32167). In a manual procedure the surface model was modified and edited in MAYA 16 (Autodesk) to create the hatchling jaw morphology and middle ear bones based on illustrations of hatchling MO 39 from Zeller (1989) and to fit the histological sections within (Fig. 2).

All 3D models (except *Ornithorhynchus*) were segmented from micro-CT-image stacks and virtual 3D models were created using Avizo 8.1 (Visualization Sciences Group, France). Editing and virtual alignment was performed with Polyworks (2014, InnovMetric Software Inc., Canada) including Gouraud filters for smoothing surfaces, adding and removing triangles, reworking mesh to reduce data size. The 3D models representing different evolutionary stages of the middle ear bones and ossified Meckel’s cartilage were manually created using MAYA 16 (Autodesk) based on illustrations in Kermack et al. (1973), Kermack et al. (1981), Luo (2011), Luo et al. (2007), Luo et al. (2016), Ji et al. (2009), and Mao et al. (2020).

## ACKNOWLEDGEMENTS

We are grateful to Thomas Martin, Rich Cifelli, Brian Davis and Pip Brewer for providing micro-CT scans of the lower jaws of *Morganucodon*, *Priacodon*, *Haldanodon* and *Dryolestes*. Sue Herring, Alec Wilken, Savannah Olroyd, James Rawson, and Abigail Tucker provided valuable feedback on earlier drafts of the manuscript. Further, we benefitted from discussions with Zhe-Xi Luo, Thomas Martin, James A. Hopson, Rebecca German, Paulina Jiménez Huidobro, Bastian Mähler, Simone Hoffman, Elsa Panciroli, Elis Newham, Pamela Gill, Callum Ross, Dallas Krentzel, Allison K. Bormet, and Kenneth-Dieter Benton. For technical support, we thank Peter Göddertz. Funding to LNW was provided by National Science Foundation (NSF) Graduate Research Fellowship and EAR-PF 2052992, and DMG was supported by an NSF Postdoctoral Research Fellowship in Biology (DBI-1812126). JAS profited from funding by the Deutsche Forschungsgemeinschaft (DFG) during the two funding periods of research unit FOR 771.

## Data availability statement

there are no data to be archived.

## Author contributions

JAS, DMG, and LNW wrote the manuscript and created the figures. JAS contributed *Haldanodon* and *Dryolestes* 3D data, creating and editing of 3D digital models. KRKJ contributed *Morganucodon* and *Priacodon* 3D data and edited the manuscript.

